# Exploring the genetic architecture of ecophysiological traits in microbial ecology using statistical analysis of the properties of protein sequences: application to the glycogen accumulating organism (GAO) phenotype

**DOI:** 10.1101/2023.10.11.561824

**Authors:** Rohan B. H. Williams, Irina Bessarab, Michael J. Wise, Krithika Arumugam

## Abstract

Nearly 50 years ago, King and Wilson introduced a key distinction between two types of genetic variation–named structural and regulatory variation–for exploring the molecular basis for phenotypic traits. Structural variation is genetic variation that will influence gene product structure via changes in coding sequence, whereas regulatory variation will influence the control of gene product expression. Here we repurpose these concepts to study the molecular basis of eco-physiological traits observed in microbial ecology, focusing on a specialized glycogen accumulating phenotype, known to be exhibited by a phylogenetically diverse group of microbes. We analyse the statistical properties of the protein sequence of the 1,4–*α*–glucan branching enzyme (*glgB*), a key enzyme responsible for formation of glycogen from linear glucans. We show that the *glgB* proteins in a subgroup of these organisms show unusual statistical properties of protein sequence length, sequence similarity and patterning of functional domains, compared to organisms that do not exhibit this phenotype. These findings suggest a role for structural genetic variation in determining this phenotype in some species. Our analysis holds implications for dissecting the genetic/genomic architecture of complex traits exhibited by microbial communities and also provides a complementary framework to presence–absence level analyses that are ubiquitous in microbial ecology.

---

The glycogen accumulating organism (GAO) phenotype is a specialized adaptation of the glycogen storage machinery, operative in cyclic anaerobic-aerobic environments, and characterised by the preferential (excessive) storage and utilisation of glycogen as both carbon and energy source. The GAO phenotype was first identified in wastewater (activated sludge) bioprocess reactors in a phylogenetically diverse set of organisms, including members of the alphaproteobacterial genus *Defluviicoccus*, the *Candidatus* Competibacter within *Gammaprotebacteria* and members of the *Actinobacteria* genera *Micropruina, Nakamurella* and *Kineosphera* [1]. Many of these species easily uptake organic substrates and rapidly convert them to storage polymers, rather than using them to support growth [2]. Given that glycogen storage is widely utilised in the prokaryotic kingdom –with recent estimates showing that around 30% of prokaryotic genomes harbour the classical pathway for glycogen biosynthesis [3]–a logical question to ask is whether the glycogen storage machinery of GAO species exhibit unusual or enhanced metabolic properties compared to those found in non-GAO species? Addressing this question will hold broader implications for microbial ecology, by providing tools for understanding how genetic (or genomic) variation influences the architecture of functional traits exhibited by microbial communities. Developing such analyses will enable an alternative functional analytic framework than the current focus on analysing the presence or absence of a given genes(s) that are currently dominant in the field.

We can frame this question using concepts introduced by King and Wilson almost half a century ago, who introduced a key distinction between two types of genetic variation– named structural and regulatory variation–for exploring the molecular basis for phenotypic traits [4]. Structural variation is genetic variation that will influence gene product structure, and potentially, function, via changes in coding sequence, whereas in the case of regulatory variation, genetic variation will influence the control of gene product expression, for example, via sequence in *cis*–regulatory control regions, which may alter a phenotypic outcome. Or in other words, is the GAO phenotype largely driven by superior metabolic capacity of the component proteins in glycogen storage pathways (the structural hypothesis) or is it a consequence of optimal regulation of those pathways in the specific environments where the phenotype is observed? (the regulatory hypothesis). To address the regulatory hypothesis, we would, at minimum, require an ensemble of culturable GAO and non–GAO species, along with assays for measuring the dynamics of glycogen formation. While some, but not all GAO species are culturable, they are typically slow growing [2, 5] and are therefore not well aligned to experimental requirements. Furthermore accurate measurements of intracellular glycogen remain challenging to obtain [6]. It would appear easier to approach the problem via the structural hypothesis: that the GAO phenotype is, or is not, a consequence of structural variation in the relevant cellular machinery. This can be done by simply examining the properties of the relevant protein sequences. We develop such an analysis here, focusing on the 1,4– alpha-glucan branching enzyme (encoded by *glgB*), which is responsible for formation of glycogen from linear glucans [7, 8, 9]. We focus on analysing protein sequence length, degree of sequence similarity, and domain composition and organisation, as the primary measures of interest.

We collated a group of 60 *glgB* protein sequences from a group of GAO species, namely genomes from eight members of genus *Defluviicoccus* (*n* =37), *Candidatus* Competibacter (*n* =6), *Candidatus* Contendobacter (*n* =3), *Propionivibrio* (*n*=2), genus *Micropruina* (*n* =1), genus *Nakamurella* (*n* =10), genus *Kinosphaera* (*n* =1) or genus Propionvibrio (*n* =2). Control groups included all prokaryotic *glgB* protein sequences from RefSeq (*n* =30,489), and two sets of ecobiological controls, firstly a subset of sequences (*n* =38) from 6 genera associated with the polyphosphate accumulating organism (PAO) phenotype: some species of which utilise glycogen in the same manner as GAO species in the anaerobic phase of anaerobic–aerobic cycling [10, 11, 12]–and secondly a set of *glgB* sequences from an activated sludge (AS) community (*n* =5012; data obtained from our previous analysis [13]).

Previous analyses of *glgB* protein sequences have noted a bimodal distribution of two characteristic sequences lengths, whose peaks are at around 640 a.a. and 730 a.a., respectively. This distribution was clearly visible in both the RefSeq and AS datasets (Fig. 1A), with the range of observed lengths (quoted as 1st-%ile to 99th-%ile) ranging from 477–1243 a.a. and 88-1219 a.a., respectively. Interestingly, we noted that in the GAO *glgB* sequences, there was an enrichment for longer forms of the protein (Fig. 1A– 1B), with 17% of GAO sequences being *>*1000 a.a. compared to no more than 1.86% in any of the control groups (*P* = 2.2 *×* 10^*−*16^; *χ*^2^ test for proportions). This enrichment was not present in *glgB* from the PAO species, whose length distribution was consistent with the expected background distribution (Fig. 1B). Examining *glgB* copy number, we also noted that GAO species had a greater number of *glgB* genes than observed in any control groups (Fig 1C), although this trend was not statistically significant (*P* = 0.11, Kolmogorov-Smirnov test between copy number distributions in RefSeq and GAO species). Longer forms of glgB were observed in some, but not all, GAO species, occurring in *Defluviicoccus, Candidatus* Competibacter, *Candidatus* Contendobacter and *Kinosphaera* (Table S1).

**Figure 1:**
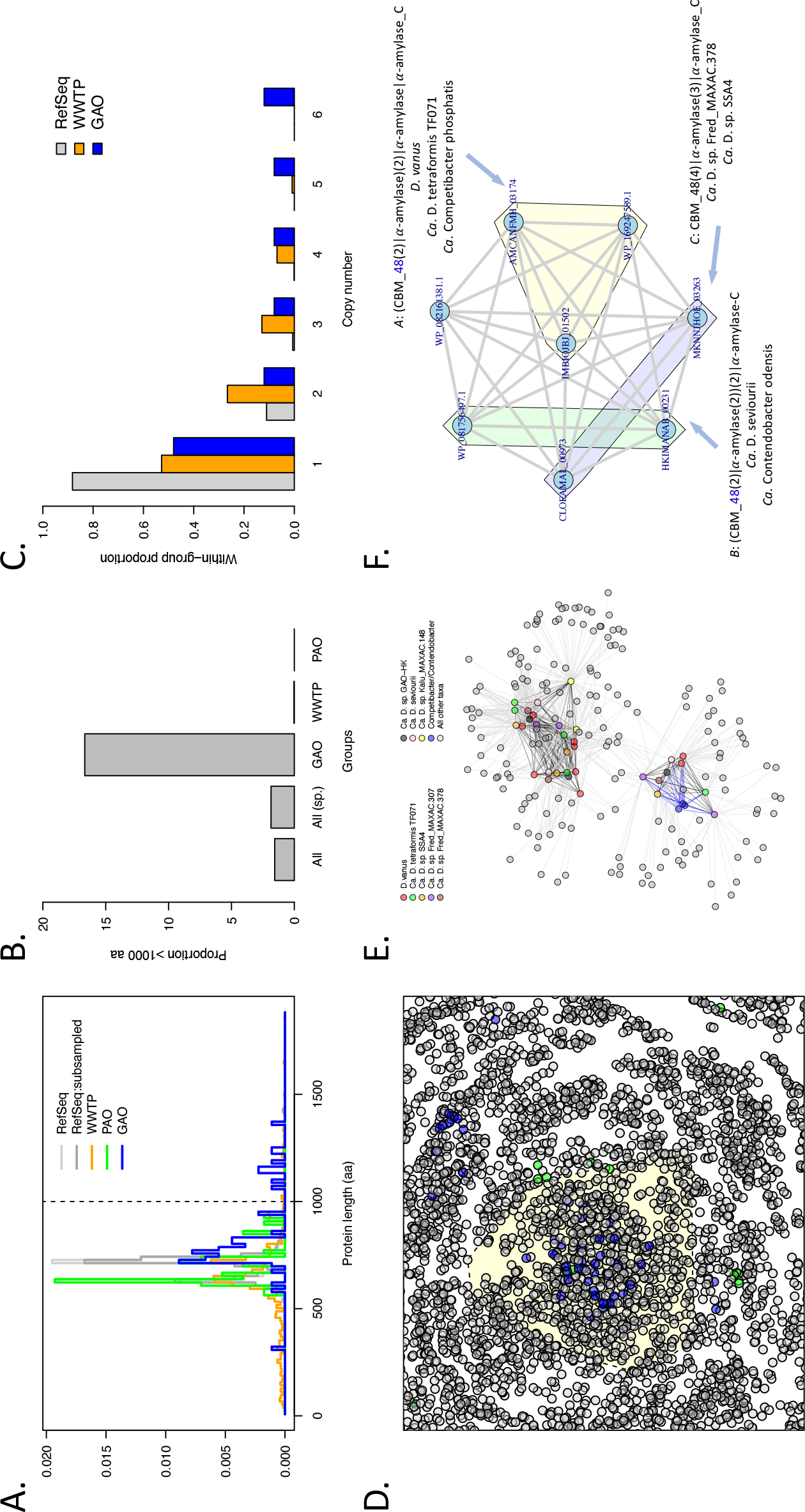
Properties of *glgB* protein sequence. *A)*: distribution of *glgB* protein sequence length in GAO species and related control populations; *B)*: proportion of *glgB* that attain a length greater than 1000 amino acids, note the enrichment for longer forms of *glgB* in GAO species (*P* = 2.2*×*10^*−*16^; *χ*^2^ Test for Proportions); *C)*: distribution of *glgB* copy number in GAO, WWTP and RefSeq groups. GAO species tend to have more copies of *glgB* encoded in their genomes but this is of only moderate statistical significance (*P* =0.11, K–S test). *D* : a portion of the main component of the *glgB* sequence similarity network, showing several regions enriched for *glgB* from GAO species (blue); *glgB* from PAO species shown in green and all others are shown in grey. Edges are not displayed for clarity. The GAO–rich region highlighted by the light yellow convex hull is expanded in *E*) and shows two distinct sub–clusters of protein sequences. The lower sub–cluster contains a clique structure that is expanded in *1F)*. Within the clique, there are several types of protein domain occurrence patterns in log forms of *glgB* that are only observed in GAO species (labelled A–C).

To examine relationships among *glgB* protein coding sequences, a sequence similarity network [14] was constructed from all *glgB* protein sequences available from NCBI. The network was comprised of 60 components, which contained between 2 and 24509 sequences (mean 509; median 50). The large main component (24,509 proteins; shortest path diameter of 19) contained 85% (51/60) of the GAO sequences. These were relatively tightly clustered, with the mean shortest path length between GAO sequences being 3.2 (range: 1–9). Sequences from GAO species were distinct from those of PAO species (*F* =23.2, *R*^2^ =0.21; *P <* 0.001; PERMANOVA between shortest path of GAO and PAO species; Fig. 1D and Fig. S1), while occupying several distinct subregions of the main component (Fig. 1E). One such region (highlighted by the yellow convex hull in Fig. 1E and expanded in Fig. 1F) contained 42 of the 51 GAO *glgB* sequences (82%), and this subset included the 10 long forms of *glgB*, 9 of which were observed to be in GAO species (the single sequence from a non–GAO species was from Streptomyces nanshensis). Nine of these long *glgB* sequence from genus *Defluviicoccus* (*n* =6), *Candidatus* Competibacter (*n* =2 species) and *Candidatus* Contendobacter (n=1) were connected in a clique, with the mean transitivity of 1.00 compared to a background mean transitivity in the entire component of 0.56 [15]. These sequences demonstrated full length alignments (mean query:sequence ratio of 1.00, range: 0.92–1.13) with a moderate degree of identity (mean percent identity 55.47, range: 49.10–80.00). We note however that this co–occurrence pattern is consistent with the strong degree of sequence length assortativity [16] in the complete network (*r* =0.79; Fig. S2), and therefore we further examined protein domain composition and organisation to gain further insight into possible inter–relationships among these sequences.

As expected, detected protein domains were dominated by the CBM 48 (carbohydrate binding), alpha–amylase and C–terminus alpha-amylase C domains, albeit with a considerable degree of diversity observed in the copy number and patterning of first two domains (full breakdown provided in Supplementary Data File 1). From the complete set of *glgB* sequences, there were 427 unique patterns of domains observed, of which 207 (48.5%) were singletons and the remainder occurred in between 2 and 12,290 genes (mean: 138, median: 5; Supplementary Data File 1). *glgB* proteins from GAO species occurred in 17 types of domain patterns, with 8 of those 17 domain patterns only being observed in GAO species (Fig S3). Four of these 8 domains were present in single protein sequences, the remaining four being present in three duos and one trio (Fig S3). Interestingly, the three latter domain patterns only occurred within the nine–sequence clique described above (Fig. 1G); with one trio containing a common domain pattern occurring in two *Defluviicoccus* species and *Candidatus* Competibacter phosphatis (Fig 2A). One duo contained a common domain pattern observable in one *Defluviicoccus* species and *Candidatus* Contendobacter odensis (Fig. 2B), while the other duo was observed between another two *Defluviicoccos* species (Fig 2C). We note that some species that share these domain patterns hold distant phylogenetic relationships, with the lowest common taxonomic level being phylum *Proteobacteria* in the NCBI taxonomy.

**Figure 2:**
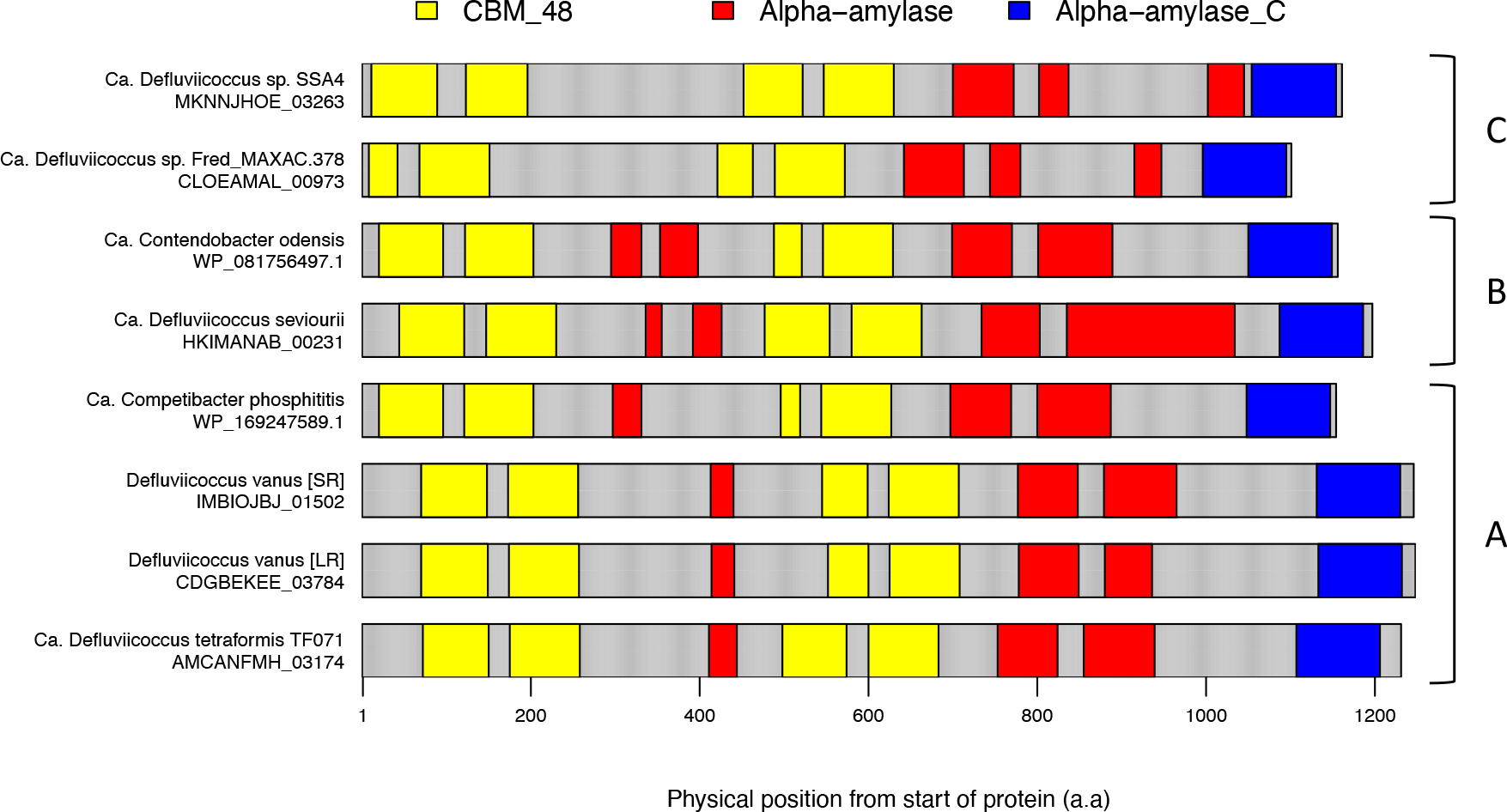
Patterns of CBM48, *α*-amylase and C–terminus *α*-amylase domains in the three sub–types of *glgB* highlighted in Fig. 1F. Letter annotations on right hand side correspond to those used in Fig 1F. For A, the cognate gene from both short– and long–read assemblies of *D. vanus* is displayed to highlight reproducibility.

Collectively, our analysis supports the view that some GAO species harbour unusual structural variants of the *glgB* enzyme: while we have no direct evidence of their functional consequences, previous lines of evidence suggest that these structural variants are likely to have significant phenotypic impact.

Previous experimental data obtained from mutant *glgB* genes clearly show that structural genetic variation will alter glycogen structure, abundance, glycogen granule morphology and cellular viability [17, 18, 19, 20]. Empirically, the average chain length of glycogen shows wide variation across the prokaryotic kingdom, and meta–analyses suggest there exists an inverse relationship between average chain length and bacterial durability, possibly as a consequence of the increased metabolic activity required to degrade short chain length glycogen [21, 22]. These data suggest that altering both the capacity for substrate binding and branch formation, as observed by variable repeats of CBM– and amylase–domains, respectively, are likely to impact glycogen formation and utilisation in the relevant GAO species. Further studies using both experimental and computational structure determination are warranted, as are expression studies with detailed biochemical and functional characterisation. Data from these non–canonical *glgB* may also have translational applications for the practical use in resource recovery and industrial application [23].

In our current understanding, GAO species are, in terms of their ecological strategy, exhibit behaviour aligned with the classical *K*–strategist categorisation [2, 24]. This behaviour can be characterised by slow growth rates, accumulation of energy reserves and robustness to environmental perturbation, with intracellular glycogen as one key energy storage system that could underpin these ecophysiological phenotypes [2, 21]. The identification of increased *glgB* copy number and an enrichment for longer forms of *glgB* suggest that some GAO may harbour an enhanced or extended machinery for storing and utilising glycogen. While our analysis shows that many other species harbour such non–canonical forms of *glgB*, we identify several functional domain patterns that are selective for phylogenetically diverse GAO species, whose closest taxonomic relationship is at phylum level. Whether these shared functional motifs arise from consequence of horizontal gene transfer or convergent evolution in these species under related selection pressures is unclear, and would require further analysis.

We have made use of the distinction between structural and regulatory genetic variation, concepts and terminology originally introduced by King and Wilson [4]. Although the presence of regulatory variation is harder to evaluate, it is likely that it will play an important role, both within and between species, in influencing variation in glycogen metabolism. We would like to highlight a recent paper by McDaniel and colleagues [25] reporting a method called TBasCO, which explores the use of molecular phenotypes (mRNA levels) in ensembles of expressed orthologous genes identified within a microbial community. Studying the dynamics of mRNA levels within these orthologous gene groups, the authors identify distinct groups of taxa, as inferred from clustered mRNA level profiles, which would logically reflect the influence of inter–specific regulatory variation. Such approaches, combined with analysis of *cis*–regulatory sequence [26, 27], may provide new insights into the influence of regulatory variation on functional phenotypes in complex microbial communities.

## Methods

To obtain protein sequences for *glgB* (1,4–*α*–glucan branching enzyme) the NCBI Protein database was searched using the term “glgb[All Fields]” with search results restricted to the bacterial kingdom within NCBI RefSeq: in total 30,489 glgB protein sequences were downloaded and augmented with 43 *glgB* sequences identified in the *Defluviicoccus* genomes obtained and/or sourced by Bessarab [28]. *glgB* from known GAO and PAO species, catalogued by Stokholm–Bjerregaard [1], were identified from taxonomic annotations. From the entire set of 30,532 protein sequences, an all–against–all homology search was conducted with DIAMOND blastp (v2.0.12.150, with default settings) [29] and the tabular output was imported into R (latest version used was 4.3.0) [30]. Self–self alignments were then removed and the best hit (highest bit–score) for unique query–subject combination were retained. The entire set of resultant alignments were treated as a node–edge list and used to construct a sequence similarity network using the R package igraph [31], which was used for all network analysis. The domain structures of all 30,352 glgB protein sequences was examined using HMMER3 [32], scanned against all 16,712 Hidden Makov Models (HMM) of the PFAM database [33]. Subsequent data analysis on the presence and duplication of specific domain was performed within the R statistical computing environment.

## Author Contributions

The concept was developed by R.B.H.W and I.B. The analysis was designed by R.B.H.W and undertaken by R.B.H.W, K.A., M.J.W and I.B. The paper was written by R.B.H.W with specific contributions and/or editing from other authors.

## Acknowledgements

This research was supported by the Singapore National Research Foundation and Ministry of Education under the Research Centre of Excellence Programme.

---

**Supplementary Table 1:**
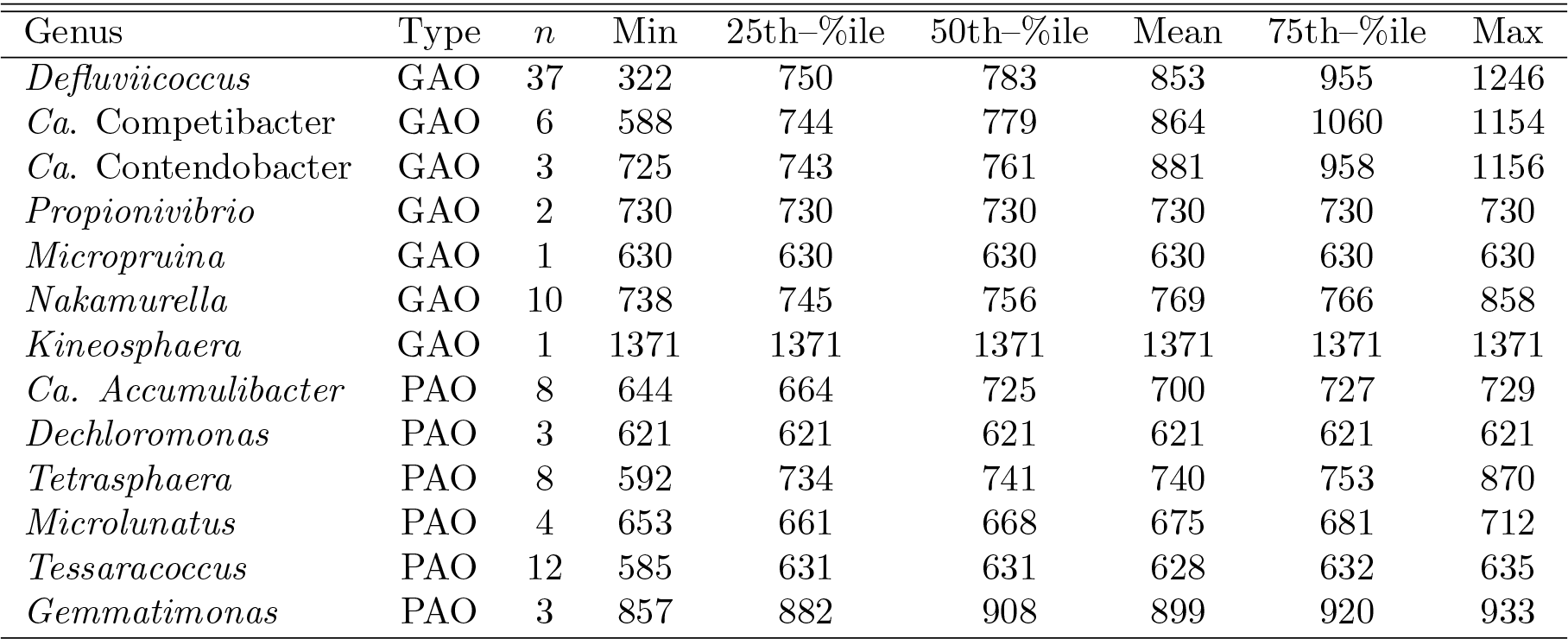
Summary statistics: *glgB* sequence length in GAO and PAO species.

**Supplementary Figure 1:**
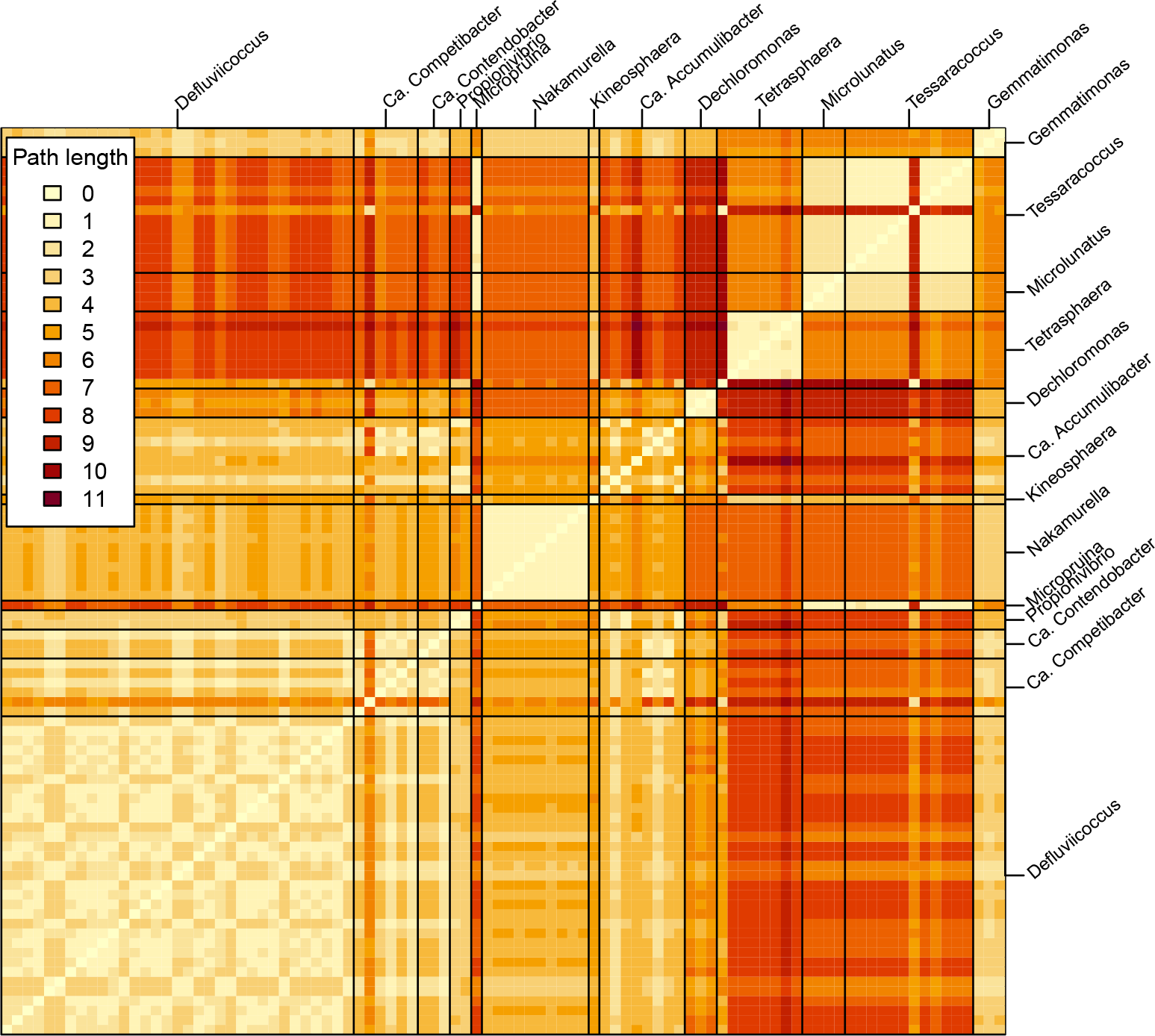
Heatmap of shortest path length in the main component of the *glgB* sequence similarity network for GAO and PAO species.

**Supplementary Figure 2:**
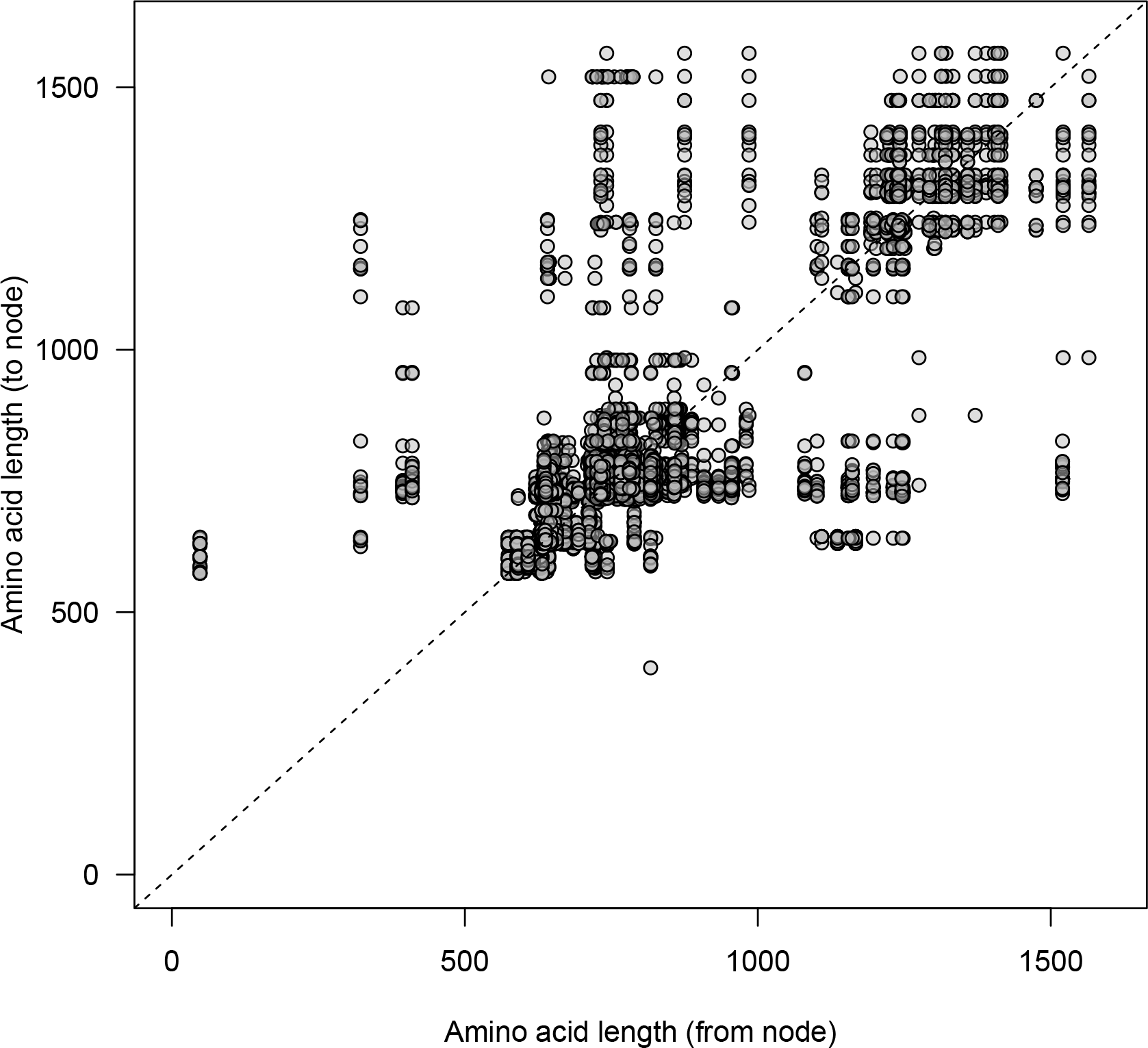
Length assortativity in the *glgB* sequence similarity network. Each data points represents an edge in the network. The *x*–axis shows the length of the *from*–node and the *y*–axis shows the length of the *to*–node. The assortativity coefficient is 0.79.

**Supplementary Figure 3:**
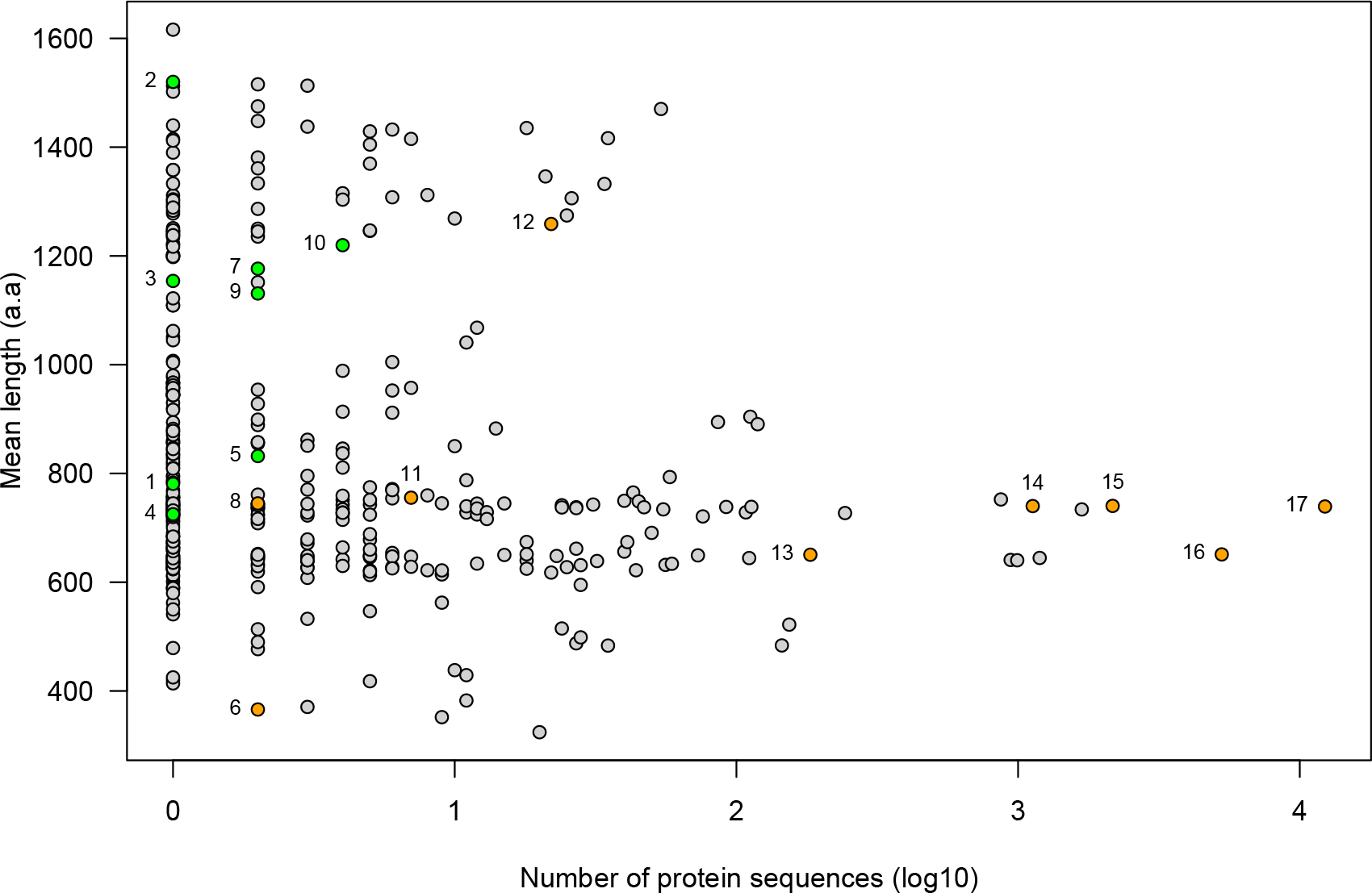
Properties of types of domain patterns observed in *glgB* sequences. Each data point represents a unique type of domain pattern observed in 30,532 *glgB* sequences. The horizontal axis shows the number of protein sequences that a domain pattern is observed in and the vertical shows their mean length (aa). Data points in yellow denote a domain pattern type that a *glgB* from a GAO-species is observed in. Data points in green denote domain pattern types that only occur in *glgB* from GAO species. For numeric annotations refer to Supplementary Data File 1. The number annotations 10, 7 and 9 correspond to the Fig. 1 letter annotations A, B and C, respectively.

## References

[1] Stokholm–Bjerregaard M, McIlroy SJ, Nierychlo M, Karst SM, Albertsen M, Nielsen PH (2017). A critical assessment of the microorganisms proposed to be important to enhanced biological phosphorus removal in full–scale wastewater treatment systems. Frontiers in Microbiology 8: 718. 10.3389/fmicb.2017.00718

[2] Seviour RJ, Maszenan AM, Soddell JA, Tandoi V, Patel BK, Kong Y, Schumann P (2000). Microbiology of the ‘G–bacteria’ in activated sludge. Environmental Microbiology 2(6): 581–93. 10.1046/j.1462-2920.2000.00153.x

[3] Wang L, Yang J, Huang Y, Liu Q, Xu Y, Piao X, Wise MJ (2019). Systematic analysis of metabolic pathway distributions of bacterial energy reserves. G3 (Bethesda) 9(8): 2489–2496. 10.1534/g3.119.400123

[4] King MC, Wilson AC (1975). Evolution at two levels in humans and chimpanzees. Science 188(4184): 107–16. 10.1126/science.1090005

[5] Maszenan AM, Bessarab I, Williams RBH, Petrovski S, Seviour RJ (2022). The phylogeny, ecology and ecophysiology of the glycogen accumulating organism (GAO) Defluviicoccus in wastewater treatment plants. Water Research 221: 118729. 10.1016/j.watres.2022.118729

[6] Wang L, Wise MJ (2019) An updated view on bacterial glycogen structure. Microbiology Australia 40: 195–199. 10.1071/MA19056

[7] Chandra G, Chater KF, Bornemann S (2011). Unexpected and widespread connections between bacterial glycogen and trehalose metabolism. Microbiology (Reading) 157(Pt 6): 1565–1572. 10.1099/mic.0.044263-0

[8] Zmasek CM, Godzik A (2014). Phylogenomic analysis of glycogen branching and debranching enzymatic duo. BMC Evolutionary Biology 14: 183. 10.1186/s12862-014-0183-2

[9] Rashid AM, Batey SF, Syson K, Koliwer–Brandl H, Miah F, Barclay JE, Findlay KC, Nartowski KP, Khimyak YZ, Kalscheuer R, Bornemann S (2016). Assembly of α–glucan by GlgE and GlgB in Mycobacteria and Streptomycetes. Biochemistry 55(23): 3270–3284. 10.1021/acs.biochem.6b00209

[10] Nielsen PH, McIlroy SJ, Albertsen M, Nierychlo M (2019). Re–evaluating the microbiology of the enhanced biological phosphorus removal process. Current Opinion in Biotechnology 57: 111–118. 10.1016/j.copbio.2019.03.008

[11] Roy S, Guanglei Q, Zuniga–Montanez R, Williams RB, Wuertz S (2021). Recent advances in understanding the ecophysiology of enhanced biological phosphorus removal. Current Opinion in Biotechnology 67: 166–174. 10.1016/j.copbio.2021.01.011

[12] Saia SM, Carrick HJ, Buda AR, Regan JM, Walter MT (2021). Critical review of polyphosphate and polyphosphate accumulating organisms for agricultural water quality management. Environmental Science and Technology 55 (5): 2722–2742. 10.1021/acs.est.0c03566

[13] Haryono MAS, Law YY, Arumugam K, Liew LC, Nguyen TQN, Drautz-Moses DI, Schuster SC, Wuertz S, Williams RBH (2022). Recovery of high quality metagenome– assembled genomes from full-scale activated sludge microbial communities in a tropical climate using longitudinal metagenome sampling. Frontiers in Microbiology 13: 869135. 10.3389/fmicb.2022.869135

[14] Atkinson HJ, Morris JH, Ferrin TE, Babbitt PC (2009). Using sequence similarity networks for visualization of relationships across diverse protein superfamilies. PLoS One 4(2): e4345. 10.1371/journal.pone.0004345

[15] Watts DJ, Strogatz SH (1998). Collective dynamics of ‘small–world’ networks. Nature 393(6684): 440–442. 10.1038/30918

[16] Newman ME (2003). Mixing patterns in networks. Physical Review E 67(2 Pt 2): 026126. 10.1103/PhysRevE.67.026126

[17] Binderup K, Mikkelsen R, Preiss J (2002). Truncation of the amino terminus of branching enzyme changes its chain transfer pattern. Archives of Biochemistry and Biophysics 397(2): 279–285. 10.1006/abbi.2001.2544

[18] Devillers CH, Piper ME, Ballicora MA, Preiss J (2003). Characterization of the branching patterns of glycogen branching enzyme truncated on the N–terminus. Archives of Biochemistry and Biophysics 418(1): 34–38. 10.1016/S0003-9861(03)00341-2. Erratum in: Arch Biochem Biophys. 2004 421(2): 290.

[19] Huang XF, Nazarian-Firouzabadi F, Vincken JP, Ji Q, Suurs LC, Visser RG, Trindade LM (2013). Expression of an engineered granule–bound Escherichia coli glycogen branching enzyme in potato results in severe morphological changes in starch granules. Plant Biotechnology Journal 11(4): 470–479. 10.1111/pbi.12033

[20] Wang L, Regina A, Butardo VM Jr, Kosar-Hashemi B, Larroque O, Kahler CM, Wise MJ (2015). Influence of in situ progressive N–terminal is still controversial truncation of glycogen branching enzyme in Escherichia coli DH5α on glycogen structure, accumulation, and bacterial viability. BMC Microbiology 15: 96. 10.1186/s12866-015-0421-9

[21] Wang L, Wise MJ. (2011) Glycogen with short average chain length enhances bacterial durability. Naturwissenschaften 98 (9): 719–729. 10.1007/s00114-011-0832-x

[22] Wang L, Wang M, Wise MJ, Liu Q, Yang T, Zhu Z, Li C, Tan X, Tang D, Wang W (2020). Recent progress in the structure of glycogen serving as a durable energy reserve in bacteria. World Journal of Microbiology and Biotechnology 36(1): 14. 10.1007/s11274-019-2795-6.

[23] Ellis RP, Cochrane MP, Dale MFB, Duffus CM, Lynn A, Morrison IM, Prentice RDM, Swanston JS, Tiller SA (1998), Starch production and industrial use. Journal of the Science of Food and Agriculture 77: 289–311.

[24] Treseder, K. K. (2023). Ecological strategies of microbes: thinking outside the triangle. Journal of Ecology 111(9): 1832–1843. 10.1111/1365-2745.14115

[25] McDaniel EA, van Steenbrugge JJM, Noguera DR, McMahon KD, Raaijmakers JM, Medema MH, Oyserman BO (2022). TbasCO: trait–based comparative ‘omics identifies ecosystem–level and niche–differentiating adaptations of an engineered microbiome. ISME Communications 2: 111 10.1038/s43705-022-00189-2

[26] Eskin E, Keich U, Gelfand MS, Pevzner PA (2003). Genome–wide analysis of bacterial promoter regions. Pacific Symposium on Biocomputing. 2003: 29–40. PMID:12603015. Full text available at http://psb.stanford.edu/psb-online/proceedings/psb03/abstracts/p29.html

[27] Barshai M, Tripto E, Orenstein Y (2020). Identifying regulatory elements via deep learning, Annual Review of Biomedical Data Science 3: 315–338. 10.1146/annurev-biodatasci-022020-021940

[28] Bessarab I, Maszenan AM, Haryono MAS, Arumugam K, Saw NMMT, Seviour RJ, Williams RBH (2022). Comparative genomics of members of the genus Defluviicoccus with insights Into their ecophysiological importance. Frontiers in Microbiology 13: 834906. 10.3389/fmicb.2022.834906.

[29] Buchfink B, Xie C, Huson DH (2015). Fast and sensitive protein alignment using DIAMOND, Nature Methods 12: 59–60. 10.1038/nmeth.3176

[30] R Core Team (2021). R: A language and environment for statistical computing. R Foundation for Statistical Computing, Vienna, Austria. https://www.R-project.org/

[31] Csrdi G, Nepusz T, Traag V, Horvt S, Zanini F, Noom D, Mller K (2023). igraph: Network analysis and visualization in R. doi:10.5281/zenodo.7682609, R package version 1.5.1, https://CRAN.R-project.org/package=igraph.

[32] S. R. Eddy (2011). Accelerated profile HMM searches. PLOS Computational Biology 7: e1002195. http://hmmer.org/

[33] El–Gebali S, Mistry J, Bateman A, Eddy SR, Luciani A, Potter SC, Qureshi M, Richardson LJ, Salazar GA, Smart A, Sonnhammer ELL, Hirsh L, Paladin L, Piovesan D, Tosatto SCE, Finn RD (2019). The Pfam protein families database in 2019. Nucleic Acids Research 47(D1): D427–D432. 10.1093/nar/gky995

